# Highly efficient DNA-free gene disruption in the agricultural pest *Ceratitis capitata* by CRISPR-Cas9 RNPs

**DOI:** 10.1101/127506

**Authors:** Angela Meccariello, Simona Maria Monti, Alessandra Romanelli, Rita Colonna, Pasquale Primo, Maria Grazia Inghilterra, Giuseppe Del Corsano, Antonio Ramaglia, Giovanni Iazzetti, Antonia Chiarore, Francesco Patti, Svenia D. Heinze, Marco Salvemini, Helen Lindsay, Elena Chiavacci, Alexa Burger, Mark D. Robinson, Christian Mosimann, Daniel Bopp, Giuseppe Saccone

**Author notes:** Author for correspondence: Giuseppe Saccone, Department of Biology, University of Naples “Federico II” Complesso Universitario Monte Sant’Angelo Building 7, room 0F-01 Via Cinthia, n. 26, 80126, Napoli, Italy.

## Abstract

The Mediterranean fruitfly *Ceratitis capitata* (medfly) is an invasive agricultural pest of high economical impact and has become an emerging model for developing new genetic control strategies as alternative to insecticides. Here, we report the successful adaptation of CRISPR-Cas9-based gene disruption in the medfly by injecting *in vitro* pre-assembled, solubilized Cas9 ribonucleoprotein complexes (RNPs) loaded with gene-specific sgRNAs into early embryos. When targeting the eye pigmentation gene *white eye* (*we*), we observed a high rate of somatic mosaicism in surviving G0 adults. Germline transmission of mutated *we* alleles by G0 animals was on average above 70%, with individual cases achieving a transmission rate of nearly 100%. We further recovered large deletions in the *we* gene when two sites were simultaneously targeted by two sgRNAs. CRISPR-Cas9 targeting of the *Ceratitis* ortholog of the *Drosophila* segmentation *paired* gene (*Ccprd*) caused segmental malformations in late embryos and in hatched larvae. Mutant phenotypes correlate with repair by non-homologous end joining (NHEJ) lesions in the two targeted genes. This simple and highly effective Cas9 RNP-based gene editing to introduce mutations in *Ceratitis capitata* will significantly advance the design and development of new effective strategies for pest control management.

## Introduction

The Mediterranean fruitfly *Ceratitis capitata* (medfly) is an economically relevant agricultural pest infesting more than 260 crop species, including fruits, vegetables, and nuts^1^. Wild populations can be contained by the Sterile Insect Technique (SIT), an eradication strategy based on the repeated release of large numbers of factory-grown sterile males into infested areas^2,3^. *C. capitata* was the first non-Drosophilidae insect species in which transposon-mediated germline transformation was established^4,5^. Various *Ceratitis* transgenic strains have been developed, aiming to improve SIT and other pest control strategies^8-16^. Also embryonic RNA interference was successfully applied to study *in vivo* functions of key *Ceratitis* genes controlling sex determination^6,7^. Nonetheless, a more comprehensive study of gene functions in *Ceratitis* will be needed to further improve existing control strategies. To generate long-lasting and hereditable changes in gene function, the CRISPR-Cas9 system with its modular and simple components provides a promising tool to implement scalable, reproducible pest control strategies^17,18^. Furthermore, transgene-based CRISPR-Cas9 can be used to produce homozygous loss-of-function mutations as well as a novel gene drive system for insect population control^19,21^.

Various studies have reported the successful use of the Cas9 system to introduce genome modifications in *Drosophila melanogaster* based on injecting different combinations CRISPR-Cas9 reagents into embryos, such as DNA plasmids expressing Cas9 protein plus single-guide RNA (sgRNA)^22^, *in vitro*-transcribed *Cas9* mRNA plus sgRNA^22,23^, and sgRNA into transgenic flies that express Cas9 in the germ line^25,26^. Lee *et al*., 2014 injected purified Cas9 protein preloaded with 2 RNAs, trRNA (transactivating RNA) and gene-specific crRNA (CRISPR RNA), into *Drosophila* embryos and observed a high rate of Cas9-induced lesions^27^. As Jinek *et al*., 2012, and Basu *et al*., 2015 reported that the use of a single sgRNA molecule, combining both trRNA and crRNA, is more effective than the trRNA/ crRNA dual system^28^, we opted for this technical improvement^29^.

Altogether, we decided on the strategy of injecting Cas9 ribonucleoprotein (RNP) complexes into insect embryos for the following reasons: 1) preloaded Cas9 RNPs should act immediately following injection; 2) RNPs potentially result in higher efficiencies compared to other approaches; 3) there are potentially less off-target events^30,31^; and, 4) Cas9 protein is more stable and robust than Cas9 mRNA, and can be generated with basic protein purification strategies ^30^.

Here, we show that injecting *in vitro* generated Cas9-sgRNA RNPs into *C. capitata* embryos is highly effective at inducing mono- and bi-allelic lesions of the targeted genes in both somatic and germline cells.

## RESULTS

### Cas9-induced somatic disruption of the *we* gene

To test the feasibility of Cas9-mediated gene disruption in *Ceratitis*, we targeted the *white eye* (*we*) gene, a locus required for eye pigmentation^4,5,32^. The *Ceratitis we* gene is the ortholog of the X-linked *white* (*w*) gene in *Drosophila^5^* and is located on the *Ceratitis* fifth chromosome^33,34^. The *we* gene is an ideal target to test Cas9-mediated disruption for the following reasons: 1) lack of eye pigmentation is an easily scored phenotype; 2) a *we*-mutant *Ceratitis* strain is available to test new loss-of-function alleles for complementation, and 3) *we* function is cell-autonomous and hence somatic mutant cells can be readily detected in the adult eye.

We used CHOPCHOP^35^ to identify potential Cas9 target sequences in *we* and to design three independent sgRNAs, *we*-g1, *we*-g2, and *we*-g3 **(Fig.1A)**. Injections with unloaded recombinant Cas9 protein caused an almost twofold lower survival rate (15-19%) compared to buffer alone injections (30%), suggesting a measurable level of toxicity of Cas9 protein in *Ceratitis* embryos **(Tab. 1)**. A slightly more pronounced toxicity of Cas9 was observed during transition from larvae to pupae (0.37.-0.49 compared to 0.84).

**Figure 1.**
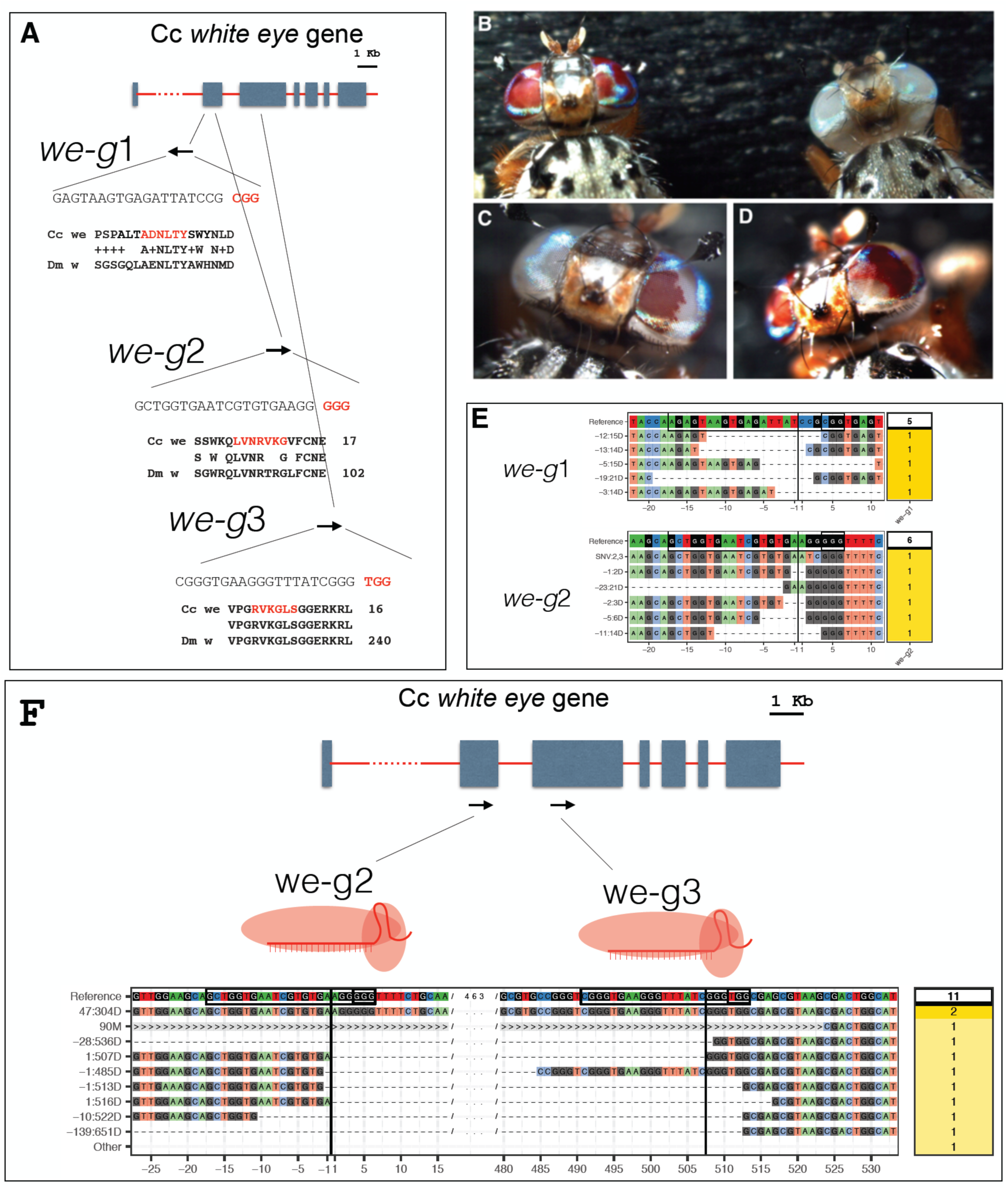
CRISPR-Cas9 targeting of *we* gene: (A) A scheme of the *we* gene and the 3 sgRNAs targeted sites (denoted *we*-g1, *we-g2, we-g3*) (B) wildtype *we*+*/we*+ (left) and mutant *we/we* individual (right). (C) and (D) examples of somatic *we* clones in the eyes of G0 individuals. (E) Sequences of mutant *we* alleles recovered from G0 individuals targeted with *we-g1* or *we*-g2. The CrispRVariants plots show sequence composition of various alleles compared to reference (target sequence and PAM are boxed and position of cut site is indicated with a black line). The label of individual allele denotes the location (relative to the cut site) and the size of the deletion. Number of sequenced clones per allele are shown in the yellow box. (F) Sequences of mutant *we* alleles in G0 individuals targeted with duplex *we-g2* and *we-g3* RNPs The CrispRVariants plot shows the spectrum of induced deletions in G0 individuals. Black lines indicate the position of the cute site in each target sequence (black boxes). > symbol signifies absence of mapped sequence from a partial alignment.

We next loaded recombinant Cas9 protein with individual sgRNAs *in vitro* and injected the RNP complexes into early syncytial embryos of the wildtype Benakeion strain. We aimed at targeting syncytial nuclei to maximize the efficiency of inducing NHEJ lesions. A biallelic hit at this early stage is expected to produce large clones of mutant tissue in injected individuals^22,30^. In this first round, we injected *we*-g1 RNPs in a buffer containing 300 KCl mM to improve solubilization of Cas9, as previously established to achieve maximal Cas9 activity in zebrafish^30,36^. Of 240 injected embryos, 64 larvae hatched and 6 survived to adulthood **(Tab. 2)**. Three displayed a mosaic pattern of *white* unpigmented ommatidia surrounded by wild-type pigmented ommatidia (Fig. 1B, C and D). One of the two eyes in one individual was completely white, suggesting that a biallelic gene disruption event occurred at an early stage in the primordial lineage (Fig. 1C). The presence of mutations was confirmed by sequencing of PCR products spanning the cleavage sites. Direct sequencing of amplified genomic DNA showed heterogeneity in nucleotide calls around the cleavage site close to the protospacer-adjacent motif (PAM) site, consistent with a range of different NJEH-induced alterations. Genomic *we* PCR products were obtained from pools of injected larvae or single adult G0 flies. Indels, mostly deletions, of variable length (Fig.1E; *we*-g1, 14-21 bp), were detected in cloned fragments, consistent with previous studies^18^.

We performed additional rounds of embryos injections with *we*-g2 (150 mM KCl), and *we*-g3 (300 mM KCl), and we subsequently observed either a lower percentage of adults with eye pigmentation mosaicism (4% for *we*-g2) or none (we-g3). Injections of *we*-g1/*we*-g2 RNPs (150 mM KCl) led to 23% adult mosaics (Tab. 2), but this duplex RNP targeting did not produce any deletions between the two targeted sites; furthermore, only *we*-g2 induced indels were observed (reported in Fig. 1E, *we*-g2: 4-6). Again, sequencing of cloned PCR products revealed NHEJ deletions, ranging from 2-21 bp, relative to the PAM site of *we*-g2. 3 additional series injections of *we*-g1+ *we*-g2 into a total of 600 embryos (data not shown), also at 300 mM KCl, confirmed the lack of deletion between the two targeted sites by PCR analysis on genomic DNA. The absence of a deletion spanning the two targeted sites less than 100 bp apart can be explained, for example, by steric hindrance which prevents that two adjacent Cas9-sgRNA complexes can cut both sites simultaneously. Indeed, when injecting simultaneously *we*-g2 and *we*-g3 RNPs (in 300 mM KCl) in early embryos, we did recover deletions spanning the 2 more distant targeted sites, ranging from 304 bp to 651 bp (Fig.1F).

### Cas9-induced germ line disruption of the *we* gene

To test for germline transmission of Cas9-induced *we* alleles, 9 injected G0 red eyed flies (3 *we*-g1/RNP flies and 6 *we*-g2 RNP flies; **Tab. 3**) were individually crossed with *we*-mutant partners (carrying a *w1* allele having a frameshift mutation in the sixth exon^32^; Genbank AH010565.2). Seven injected flies sired G1 progeny, six of which gave rise to mutant *white eye* flies (Fig.2A), with a highly variable transmission rate (1.5% - 100%; **Tab. 3**). Non-complementation of the CRISPR-Cas9-induced mutations confirms that they are allelic to the original *we* mutation (Fig. 2A). Of the three *we*-g1 injected individuals, two males sired small batches of progeny in which 100% (*we*-g1#1; 6 flies) and 45% (*we*-g1#2; *10 out of 22*), respectively, displayed the mutant phenotype (Tab. 3). Of the six *we*-g2 injected individuals, 4 males produced various proportions of G1 mutant white-eyed progeny. Remarkably, the *we*-g1#1 and *we*-g2#5 lines gave rise to 100% and 98% G1 white-eyed offspring, respectively (Tab. 3). Thus, our results demonstrate that Cas9 activity is highly effective in the germ line, producing mostly mutant primordial germ cells.

**Figure 2.**
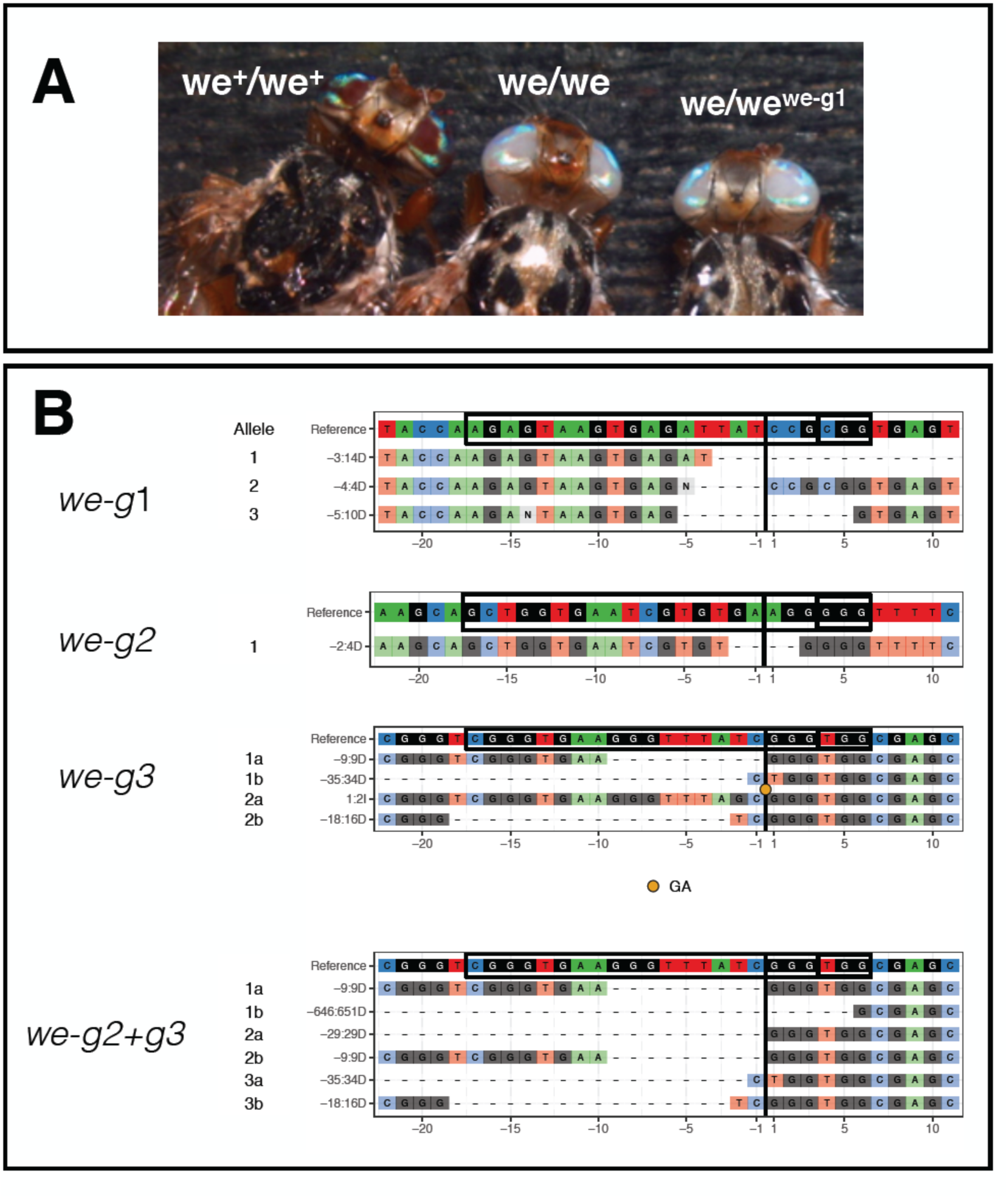
CRISPR induced *we* non-complementing alleles transmitted to the G1 progeny. (A) homozygous wildtype *we*^+^ individual (left), homozygous mutant individual *we* (middle), G1 individual heterozygous for mutant *we* and CRISPr induced *we* mutation. (B) CrispRVariants plots of G1 mutant progeny of G0 targeted by *we-*g1 (3 sequences), targeted by *we*-g2 (1 sequence), *we-g3* (6 sequences) or *we*-g2+g3 (4 sequences).

We randomly chose 4 out of 16 G1 *we*-g1-targeted mutant flies (two from *we*-g1#1 cross and two from *we*-g1#2 cross) and 2 out of 166 *we*-g2-targeted mutant G1 flies (one from *we*-g2#4 line and one from *we*-g2#5 line)(Tab. 3). Sequencing of cloned PCR products revealed that the 4 *we*-g1 G1 flies inherited one allele from the *we* parent and three different *we^CRISPR^* alleles from the injected G0 fathers. We found an identical *we* deletion of 14 bp (labeled “-3:14D” in Fig.1E and Fig.2B) in both analyzed flies from the *we-*g1#2 cross, suggesting that they inherited the same mutation from their common male founder (Fig.2B: *we*-*g1*). Mutant G1 flies of the *we*-g1#2 line carried two different alleles (4 bp and 10 bp deletions; -4:4D and -5:10D, respectively). All three Cas9-induced alleles are small deletions, causing frame-shifts in exon 2 of the *we* coding region, and failed to complement the original *we* mutation. Two *we*-g2 targeted G1 mutant flies carried two novel alleles, one with a 4 bp deletion (we-g2#4 line; Table 3) and one with a long 84 bp deletion (we-g2#5 line; Table 3) (Fig.2B: we-*g2*; 84bp deletion not shown).

To study genes for which no mutant alleles are available, it would be useful to screen for mutant phenotypes by *inter se* crossings of G0 individuals in which the germ line has been targeted by CRISPR-Cas9. To test this possibility, we injected *we*-g3 or *we*-g2+g3 RNPs (Tab. 3; 4^th^ and 5^th^ rows). All of the surviving G0 flies, 20 and 38 respectively, did not show somatic mosaicism in the eyes (Tab. 3). When crossed *inter se* **(Tab. 4),** we recovered two mutant individuals out of 26 G1 flies (8%) from the *we*-g3 cross and three out of 184 flies (2%) from the *we*-g2+*we*-g3 cross, demonstrating the feasibility of this crossing strategy.

Sequencing of the targeted regions in 2 G1 flies of the *we-g3* cross identified, as expected, 4 novel *we* alleles. 2 out of 3 tested G1 flies of the *we-g2*+*we-g3* cross all exhibit identical 9bp deletion *we* allele at *we*-g3, most likely derived from the same G0 injected parent (Fig. 2B: we-*g3*). One of these 3 flies carried a large deletion of 651 bp resulting from duplex targeting with *we*-g2+*we*-g3 (Fig.2B: we-*g2*+*g3*).

Taken together, our analysis revealed that a cross of 20 and 38 adult G0 flies from Cas9 RNP-injected embryos can lead to 2-8% of mutant G1 flies bearing heteroallelic loss-of-function combinations within two medfly generations (less than 2 months).

### Cas9-induced somatic disruption of the *Ceratitis paired* gene

The zygotic activity of the *Drosophila paired* gene (*prd*) is required for proper segmentation of the developing embryo^37^. *Drosophila* embryos homozygous for *prd* loss-of-function alleles lack every other segment and die before hatching (pair-rule phenotype)^37^. Early segmentation events are well conserved amongst higher dipterans^38^; we therefore hypothesized that the *prd* ortholog in *Ceratitis* (Genbank JAB89073.1), will display comparable segmental defects and lethality when disrupted. We used CHOPCHOP to select one target sequences in *Ccprd* (*Ccprd*-g1) in a region encoding the conserved PRD domain (green box; Fig.3A). Two injections series with *Ccprd-g1* (300 mM KCl), resulted in survival rates of hatching embryos of 35% and 24%, respectively (Tab. 5).–A slightly more pronounced lethality of *Ccprd-*g1 was observed (0.24-0.35) compared with the *we*-g1 and *we*-g2 RNPs effects (0.43-0.44; Table 2). Comparing these values to the 33% of survivors injected with buffer alone, RNP-mediated targeting of *Ccprd* led to a 3-6 times higher mortality. Approximately 1-5% of the injected embryos showed either delayed development (late hatching rate) or arrested at late stages of embryogenesis, both effects we did not observe by injection of Cas9 alone (data not shown). Some lethality was observed at larval stages, with few individuals showing impaired locomotor activity and abnormal cuticular morphology. Only 5-11% of injected individuals developed to adulthood, with a delay of 1-2 days compared to flies injected with buffer alone or with *white* targeting RNPs (Tab. 5).

**Figure 3.**
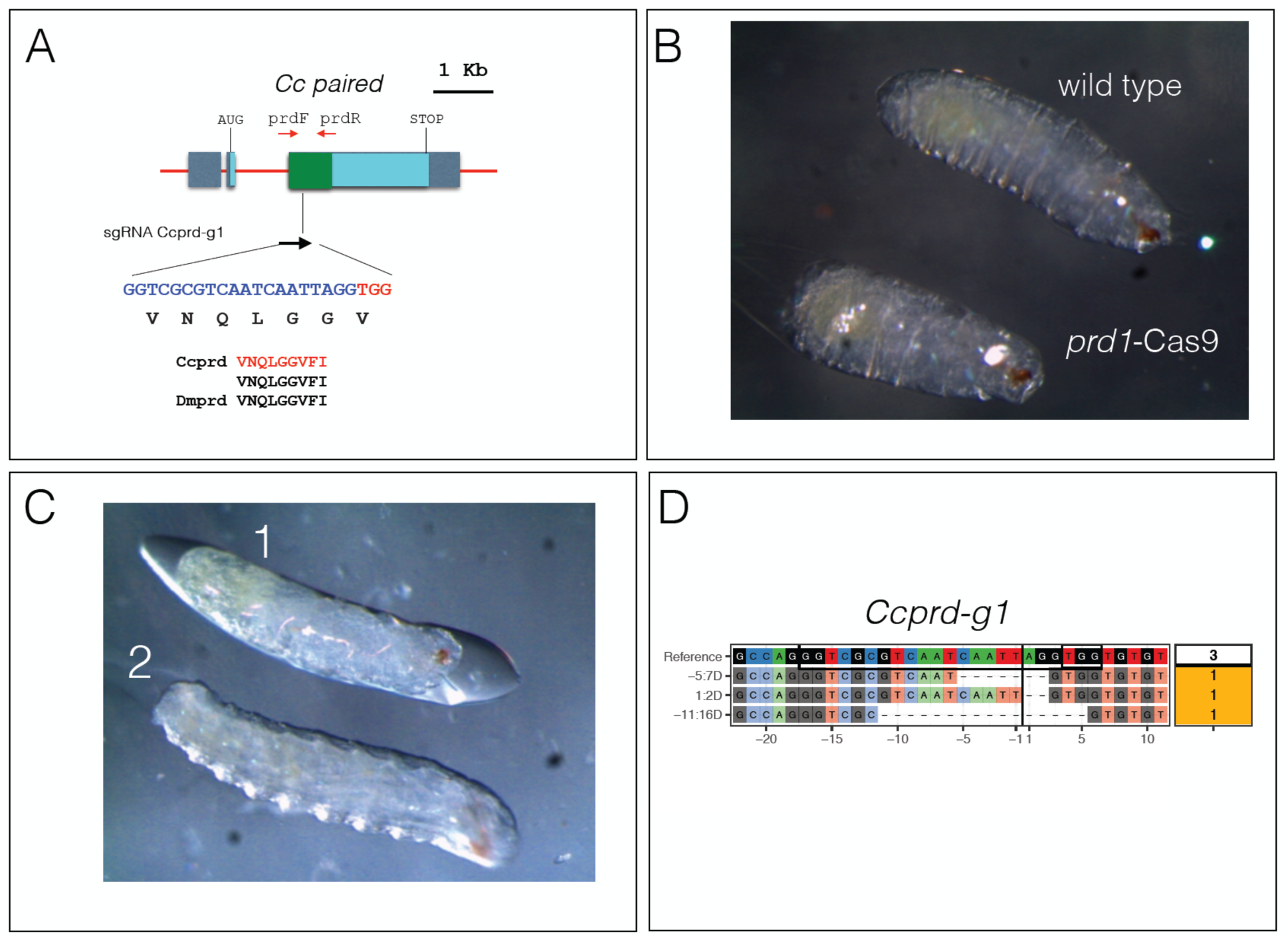
CRISP-Cas9 targeting of *Ccprd*. (A) A scheme of the *Ccprd* gene and the positions and sequences of two sgRNAs used for inducing NJEH lesions (B) Comparison of the cuticular morphology of a non-injected control (wild type) and a sgRNA prd1 injected embryo (*prd1*-Cas9). The injected embryo is significantly shorter and displays a twofold reduction in segment number reminiscent of the pair-rule phenotype described in Drosophila. (C) After injection of *Ccprd*-g1, high variability in phenotypes was observed ranging from wildtype appearance (2) to strong deformations and size reductions along the main body axis. Similar phenotypes were observed also with *Ccprd-g2* (D) A CrispRVariants plot of *Ccprd* alleles found in larvae injected with RNP containing *Ccprd-g1* sgRNA.

Approximately 90% of the embryos that failed to hatch (respectively, 90 embryos and 160 embryos; Tab. 5) displayed disorganized cuticular structures. Some were up to 50% shorter in size compared to control embryos and consisted of a reduced number of segments (Fig.3 B). This phenotype is reminiscent of the *pair rule* phenotype described for *Drosophila*^37,39^.

The phenotypic variability is likely a result of mutagenesis-caused mosaicism, with a variable proportion of wildtype cells (*Ccprd*^+^*/Ccprd*^+^ or *Ccprd*^+^*/Ccprd*^−^) and mutant cells (*Ccprd^−^/Ccprd*^−^) in these affected individuals. Because of the mosaic nature of Cas9-induced lesions in the cell-autonomously acting *Ccprd*, a complete pair rule phenotype was not expected. The observed phenotypes are consistent with the occurrence of large mutant clones caused by biallelic targeting. Sequencing of individuals injected with *Ccprd-*g1 unveiled lesions in the targeted regions of the *Ccprd* locus (Fig.3D). We conclude that *Ccprd-g1* targeting was effective in inducing lesions in *Ccprd* and causing segmental malformations.

## DISCUSSION

Over the last two decades, novel genetic strategies in pest insect management have been developed to improve their effectiveness in the field. Genetic technologies used thus far in the medfly have been based on the random integration of transposable elements into the genome^4^, site-specific modification of the randomly integrated transgene^12^ and embryonic or transgene-mediated RNA interference (RNAi)^6,15^. The disadvantage is that such genetically modified medfly must be continuously tested with respect to fitness and competitiveness as well as to stability and expression of the transgene^40^. The CRISPR-Cas9 technology offers the possibility to avoid random integration of exogenous DNA but instead provides a more robust and controlled approach to introduce new genetic features, which can be either addition of exogenous DNA or nucleotides changes in specific genes. Here, we explored the use of Cas9-sgRNA RNP complexes, avoiding plasmid-based or DNA-mediated delivery of Cas9. This DNA-free (plasmid or integrated transgene) method may allow to circumvent existing regulatory restrictions which are in effect in most nations concerning the use of GM organisms in the field. This “green” editing technology may facilitate global acceptance not only for plants or fungi, but also for insect pest control^41^.

The use of preloaded Cas9-sgRNA complexes is becoming a successful approach for targeted disruption of genes in a growing number of insect species, including *Drosophila melanogaster^27^* and *Aedes aegypti^29^*. To our knowledge, this is the first report showing that the same approach can be used to effectively mutate genes in a major agricultural pest insect such as *Ceratitis capitata*, with up to 100% mutagenesis rate in the germ line. As observed by Lee *et al*. (2014), Cas9-sgRNA complexes act immediately but are rapidly degraded, often within few hours after administration^27^; similarly Kim *et al*. (2014) reported that Cas9 protein is degraded within 24 h after being applied to cultured human cell lines^17^. As the short-lived activity of Cas9 prevents the induction of late mutational events, this may help to reduce off-target effects and mosaicism of mutant and wild type tissues in the injected individuals.

We report here that Cas9-mediated NJEH events generates lesions in the *Ceratitis we* and *Ccprd* genes. Targeting the *we* gene caused red-white eye mosaicism in up to 50% of injected individuals, indicating a very high rate of somatic bi-allelic hits. Transmission of *we* mutated alleles to G1 progeny was found to be highly effective being close to 100%. Lee *et al*., (2014)^27^ observed a 5-time lower germline transmission rate (20%) when targeting *Drosophila* genes using RNPs. One possible explanation is that the higher concentration of KCl (300 mM) used in our study increases Cas9 stability and/or solubility^30^. Injecting RNP complexes in *Aedes aegypti*, Basu *et al*., (2015) found a germline transmission rate of induced mutations up to 90%^28^; yet this rate was determined with high-resolution melt analysis (HRMA) which detects also monoallelic somatic gene editing. Instead in our study, the germline transmission rate was calculated scoring recessive G1 phenotypes, which is an under-estimation compared with HRMA. Gilles *et al*., (2015) observed up to 80% of mutant G0 progeny by co-injecting Cas9 mRNA and sgRNAs into the coleopteran *Tribolium castaneum* embryos^43^. Here, the authors scored a phenotype (lack of fluorescence) targeting a single copy dominant marker transgene. As we scored biallelic mutations of the endogenous *we* gene in G0 somatic mosaics, our efficiency rate of somatic mutation is much higher.

Several studies have reported that the simultaneous use of 2 Cas9-sgRNAs is an effective means to generate deletions of sequences between two targeted sites^29,44^. While *we*-g1 and *we*-g2 RNPs were individually effective in gene editing, the absence of a deletion spanning the two targeted sites less than 100 bp apart in 4 different series of experiments (at 150 mM and 300 mM KCl) can be explained for example by steric hindrance which prevents that two adjacent Cas9-sgRNA complexes can cut both sites simultaneously. In contrast, RNPs targeting (at 300 mM KCl) two sites more distant of each other e.g. *we*-g2 and *we*-g3 (489 bp) simultaneously was effective in generating long deletions in both somatic and germ-line cells. Hence it is conceivable that a multiplex CRISPR-Cas9 system can be used to remove for instance protein domains encoded by single exons in the medfly.

To test the functionality of purified Cas9 protein we established a fast *in vivo* assay by targeting the *Ceratitis* zygotic segmentation gene *Ccprd*. Indeed, this test allows to screen for mutant phenotypes within 3 days and subsequent sequence analysis of the targeted site can be performed to confirm the presence of lesions.

CRISPR-Cas9 will be helpful in investigating natural traits of this major agricultural pest, for example invasiveness and host adaptation, reproduction, olfaction (fruit seeking behavior), chemoreception, toxin and insecticide metabolism^45^. The recent availability of a medfly genome draft combined with the successful implementation of CRISPR genome editing technology, as reported here, opens the road to transfer basic knowledge to applied research. Next challenges for the CRISPR-Cas9 technology in *Ceratitis* will be to exploit homology-directed recombination for genome editing, either to insert transgenes in specific regions or to replace DNA sequences with slightly mutated ones.

## Methods

### Rearing of *Ceratitis capitata* and strains used

The *C. capitata* strains were reared in standard laboratory conditions at 25 °C, 70% relative humidity and 12h /12h light–dark cycles. Adult flies were fed with a mixture of yeast and sucrose powder (1v/2v). Eggs were collected in dishes filled with water and after hatching transferred to larval food (30 g, sugar 30 g, yeast extract 30 g, cholesterol stock 10 ml, HCl stock 2 ml, Benzoic stock 8,5 ml, water 400 ml). Pupae were collected and stored in dishes until eclosion.

Following strains were used in this study: a) the wildtype strain Benakeion which was obtained from P. A. Mourikis (Benakeion Institute of Phytopathology, Athens, Greece) and b) the mutant *we/we* strain Benakeion^4^ kindly provided by Kostas Bourtzis (Pest Control of FAO/IAEA, Seiberdorf, Austria).

### Cas9 protein purification

We produced our own supply of Cas9 endonuclease by expressing HIS tagged protein in bacteria^27^ and following the purification protocol described in Monti *et al*.^46^ and Dathan *et al*.^47^. The *pET* plasmid which carries the recombinant His-tagged Cas9 gene was transformed into BL21(DE3) bacterial strain. Protein expression was induced in presence of 0.5 mM isopropyl-β-D-thiogalactopyranoside (IPTG) for 16 h at 22 °C. 100 ml cell pellet was resuspended in 10 ml of cold lysis buffer (50 mM Phosphate, 500 mM NaCl, 10 mM Imidazole, 1 mM DTT, pH 8) supplemented with protease inhibitor mixture (1 mM PMSF and 1.0 mg/ml of lysozyme) and incubated at room temperature for 30 min. Cells were disrupted by sonication on ice with 10 s on and 10 s off cycles for 10 min. After centrifugation at 14,000 rpm for 30 min at 4 °C, supernatant was loaded on an ÅKTA FPLC chromatography column using a 1 ml HisTrap HP. The column was washed with lysis buffer and bound protein was eluted using a gradient from 10 mM to 500 mM imidazole. Protein elution was monitored by measuring absorbance at 280 nm and resulting fractions were analyzed by 15% SDS-PAGE. The eluted fractions were dialyzed against buffer (20 mM HEPES, 150 mM KCl, 1 mM DTT, 10% glycerol, pH 7.5). Analysis of the protein by gel-filtration was compared with calibration obtained with marker proteins run on the column under the same conditions. Gel-filtration analyses were carried out on a Superdex 200 increase 10/300 GL column.

### sgRNA synthesis and RNP complex assembly

sgRNA were designed using CHOPCHOP^35^ https://chopchop.rc.fas.harvard.edu/. CHOPCHOP lists the Target Sequence (including the PAM), the genomic location of the target, the strand (− or +), the GC content of the guide and the Off-targets. Templates for sgRNA production were designed as described by Bassett et al (2013)^23^. Following 3 templates for sgRNA production were generated to target the *we* gene:

g1:

5’GAAATTAATACGACTCACTATA<u>GAGTAAGTGAGATTATCCG</u>GTTTTAGAGCTAGAAATAGC3’

g2:

5’GAAATTAATACGACTCACTATA<u>GCTGGTGAATCGTGTGAAGG</u>GTTTTAGAGCTAGAAATAGC3’

g3:

5’GAAATTAATACGACTCACTATA<u>CGGGTGAAGGGTTTATCGGG</u>GTTTTAGAGCTAGAAATAGC3’

(respectively with PAMs CGG, GGG and TGG).

Common sgReverse (PAGE purified):

5’AAAAGCACCGACTCGGTGCCACTTTTTCAAGTTGATGGACTAGCCTTATTTTAACTTGCTATTTCTAGCTCTAAAACAAC-3’

Following 2 templates for sgRNA production were generated to target the *Ccprd* gene: Prd-g1:

5’GAAATTAATACGACTCACTATA<u>GGTCGCGTCAATCAATTAGG</u>GTTTTAGAGCTAGAAATAGC-3’

Prd-g2:

5’GAAATTAATACGACTCACTATA<u>AGAATCCCAGCATATTTTCG</u>GTTTTAGAGCTAGAAATAGC3

(respectively with PAMs TGG and TGG).

Common sgReverse (PAGE purified):

5’AAAAGCACCGACTCGGTGCCACTTTTTCAAGTTGATGGACTAGCCTTATTTTAACTTGCTATTTCTAGCTCTAAAACAAC-3’

The PCR was performed using Q5^®^ High-Fidelity DNA Polymerase (NEB). The DNA template was cleaned by phenol/chloroform extractions and EtOH precipitation at -20°C. Following 14,000 rpm 15 min 4°C, the pellet was resuspended in 40μl ddH2O. For synthesis of sgRNA we followed the instructions of Megatranscript T7 kit (Ambion) using 400 ng of target template with 5' flanking T7 promoter as starting material. After RNA synthesis, DNA template was removed by incubating with TurboDNase (Mmessage mMACHINE T7 Ultra Kit, Ambion) for 15 min at 37 °C.

The reaction for complex formation was prepared by mixing 1.8 μg of purified Cas9 protein with approximately 1 μg of sgRNA in a 5 μl volume containing 150 or 300 mM KCl, according to the protocols proposed by Lee et al. ^27^, and Burger et al.^30^, respectively. The freshly prepared mixture was incubated for 10 min at 37°C and kept on ice prior to injections.

### Microinjection of sgRNA-Cas9 RNP complexes

Embryos of the wildtype Benakeion strain were collected 1 hour after egg lay and chorion membrane was manually removed. Dechorionated embryos were aligned and glued to 3MM doubleside tape on a cover slip with posterior ends pointing to injection site and covered with Oil S700 (Sigma). A glass needle was pulled with a Narishige PB-7 and manually broken to produce sharp edges at the injection end. The needle was filled with 1 μl of preloaded sgRNA-Cas9 mix and injected into the posterior end of embryos. A Leica inverted microscope DM-IRB was used for micro-injections. After injection excess of oil was carefully removed and cover slip were put on an agar plate overnight at 25°C. Surviving larvae were transferred after 24 hours to petri dishes containing medfly larval food. A Jove video is available, describing a similar method of medfly embryos microinjection^50^

### Genomic analysis of Cas9-mediated lesions

DNA extraction was performed, with minor modifications, according to the protocol of Holmes and Bonner **[1973]**^48^. Individual larvae or adult flies were placed in a 1.5 ml Eppendorf tube and manually crushed with a pestle in 200 μl Holmes Bonner buffer (urea 7 M, Tris-HCl 100 mM pH 8.0, EDTA 10 mM pH 8.0, NaCl 350 mM, SDS 2%). Subsequently, DNA was purified by phenol/chloroform extraction, followed by chloroform extraction and ethanol precipitation. The pellet was resuspended in 30 μl or 100 μl water containing RNaseA. The resulting DNA was used as template to amplify the region encompassing the target sites, using the following primers

F-*we*: 5’GCCCTACGAGCAATCCTCT 3’

R1-*we*: 5’TCTGCAATGAGCGTCATATAC 3’

R2-*we*: 5’TTCTGCGATAGCTTTTTCAACA 3’

*F-Ccprd* 5’CTTCGACACACAACCGTGTG 3’

R-*Ccprd* 5’AGAATGCTTGTGGGAATGTTCT 3’

DreamTaq (Life Technologies) polymerase was used for PCR amplifications according to the manufacturer’s instructions. The PCR products were purified with StrataPrep PCR Purificaton Kit (Agilent Technologies) and subcloned using with StrataClone PCR cloning Kit (Agilent Techologies). Positive clones were sequenced by Sanger method and ABI 310 Automated Sequencer (Applied Biosystems). CrispRVariants and panel plots were compiled from primary Sanger sequencing data as described in ^30,49^, as well as using new functionality to display duplex guide pairs (e.g., Fig. 1F) as implemented in the development version 1.3.6.

## Acknowledgments

We wish to thank Jin-Soo Kim, Kate O’Connor-Gilles and Andrew R. Bassett for sharing information and reagents. We thank Dave O’Brochta, Max Scott and Bill Reid for invaluable help and advice and Tessa G. Montague for including the Medfly genome in the CHOPCHOP application. We wish to thank Claudia Brunner and Domenica Ippolito for technical assistance. We thank Marcelo Jacobs Lorena and Philippos Papathanos for critical reading of the manuscript. We wish to acknowledge the participation of AM in a practical course for application of the CRISPR/CAS9 technology in insects organized by Insect Genetic Technology Research Coordination Network (http://igtrcn.org; 17-21 August 2015). This project was supported by “Ricerca Dipartimentale” grants (2015, 2016), by “Legge 5/Regione Campania” grant (2008) and by a “BIP/POR Campania FESR 2007/2013” grant to GS; AM was supported with a PhD fellowship by Biology Department (University Federico II of Naples); CM received support from the Canton of Zürich, a Swiss National Science Foundation (SNSF) professorship [PP00P3_139093], and a Marie Curie Career Integration Grant from the European Commission [CIG PCIG14-GA-2013-631984]; AB was supported by an UZH URPP “Translational Cancer Research” grant; EC was supported by a project grant from the Swiss Heart Foundation.

## Contributions

AM and GS designed the experiments. AM and SMM expressed and purified the CAS9, sgRNAs design and selection was performed by AM and MR, AM produced sgRNAs, MR and HL analyzed indels and produced CrispRVariants plots. DB hosted AM and GS at University of Zurich to establish CRISPR/CAS9 technology in the medfly and in *Musca domestica;* the purified CAS9 was tested in zebrafish by CM who together with AB provided protocols and technical advices to improve Cas9 stability/efficiency; GS wrote the manuscript with major inputs from DB and contributions by AM, MR and CM. AM and GS supervised four master students, RC, MGI, PP, GDC, who contributed to this study with medfly rearing, crossings, PCR analysis, cloning and sequencing. EC, MS, FP and AC analyzed DNA clones from injected individuals. AR and GI assisted with micro-injections and video recordings.

## Additional information

**Supplementary Tables 1- 5**

## Supplementary Tables

**Supplementary Table 1:** Survival rates after injecting recombinant Cas9 protein into embryos. Numbers represent the percentage of survivors (actual number in parentheses) for a given specific developmental stage relative to number of individuals of the preceding stage.

| Cas9<br>µg/µl | Injected<br>embryos | Larvae/<br>Embryos | Pupae/<br>Larvae | Adults/<br>Pupae |
| --- | --- | --- | --- | --- |
| 0 | 114 | 0.56 (64) | 0.84 (54) | 0.66 (36) |
| 0.9 | 150 | 0.59 (88) | 0.37 (33) | 0.82 (27) |
| 1.8 | 110 | 0.57 (63) | 0.49 (31) | 0.68 (31) |
| 3.6 | 134 | 0.51 (69) | 0.41 (28) | 0.71 (20) |

**Supplementary Table 2:**
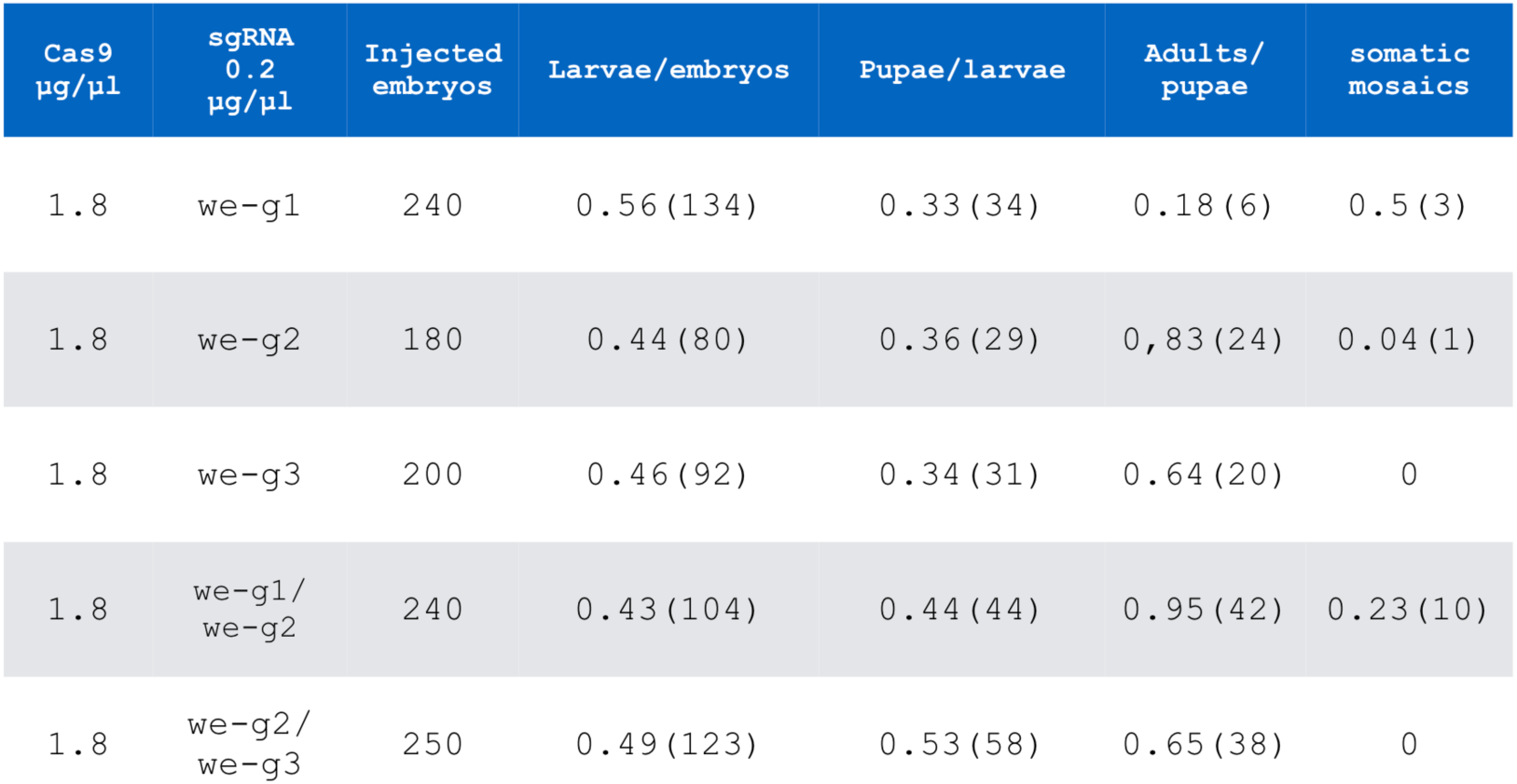
Survival rates and frequencies of individuals with somatic mutant clones after injecting we-g RNPs. Numbers represent the percentage of survivors (actual number in parentheses) for a given specific developmental stage relative to number of individuals of the preceeding stage. 31 larvae which survived *we*-g1 injections (row 1), 5 larvae of *we*-g1/*we*-g2 (row 4) and 13 larvae of *we*-g2+ *we*-g3 injections (row 5) were sacrificed for molecular analysis.

| Cas9<br>µg/µl | sgRNA<br>0.2<br>µg/µl | Injected<br>embryos | Larvae/embryos | Pupae/larvae | Adults/<br>pupae | somatic<br>mosaics |
| --- | --- | --- | --- | --- | --- | --- |
| 1.8 | we-g1 | 240 | 0.56 (134) | 0.33 (34) | 0.18 (6) | 0.5 (3) |
| 1.8 | we-g2 | 180 | 0.44 (80) | 0.36 (29) | 0.83 (24) | 0.04 (1) |
| 1.8 | we-g3 | 200 | 0.46 (92) | 0.34 (31) | 0.64 (20) | 0 |
| 1.8 | we-g1/<br>we-g2 | 240 | 0.43 (104) | 0.44 (44) | 0.95 (42) | 0.23 (10) |
| 1.8 | we-g2/<br>we-g3 | 250 | 0.49 (123) | 0.53 (58) | 0.65 (38) | 0 |

**Supplementary Table 3:** Transmission of CRISPR-induced *we* mutations in the germ line. From 9 injected G0 flies 7 (RNPs *we*-g1 or *we*-g2; see first colown) produced progeny with we phenotype when crossed to *we/we* mutant partners. 2 flies were sterile. The highest transmission rates of CRISPr induced mutant we alleles (100% and 98%) are shown in bold.

| Line | Sex of founder | *we/we*<sup>CRISPR</sup> flies | *we/we*<sup>+</sup> flies | Transmission rate |
| --- | --- | --- | --- | --- |
| *we-g1*#1 | M* | 6 | 0 | **1** |
| *we-g1*#2 | M* | 10 | 12 | 0.45 |
| *we-g2*#3 | F | 0 | 55 | 0 |
| *we-g2*#4 | M | 53 | 34 | 0.6 |
| *we-g2*#5 | M | 113 | 2 | **0.98** |
| *we-g2*#6 | M | 1 | 68 | 0.01 |
| *we-g2*#7 | M* | 14 | 10 | 0.58 |

**Supplementary Table 4:** Germline transmission of CRISPR-induced *we* mutations. Progeny of two families of G0 injected adults are shown. All G0 injected individuals from each of the 2 injection experiments shown in Table 4, were crossed and G1 scored for we mutant flies.

| sgRNAs | G0 injected parents | *we*<sup>CRISPR</sup> flies | *we*<sup>+</sup> flies | Total adult G1 flies | Mutants frequency |
| --- | --- | --- | --- | --- | --- |
| *we-g3* | 8M x 12F | 2 | 24 | 26 | 0.08 |
| *we-g2+*<br>*we-g3* | 18M x 20F | 3 | 181 | 184 | 0.02 |

**Supplementary Table 5.**
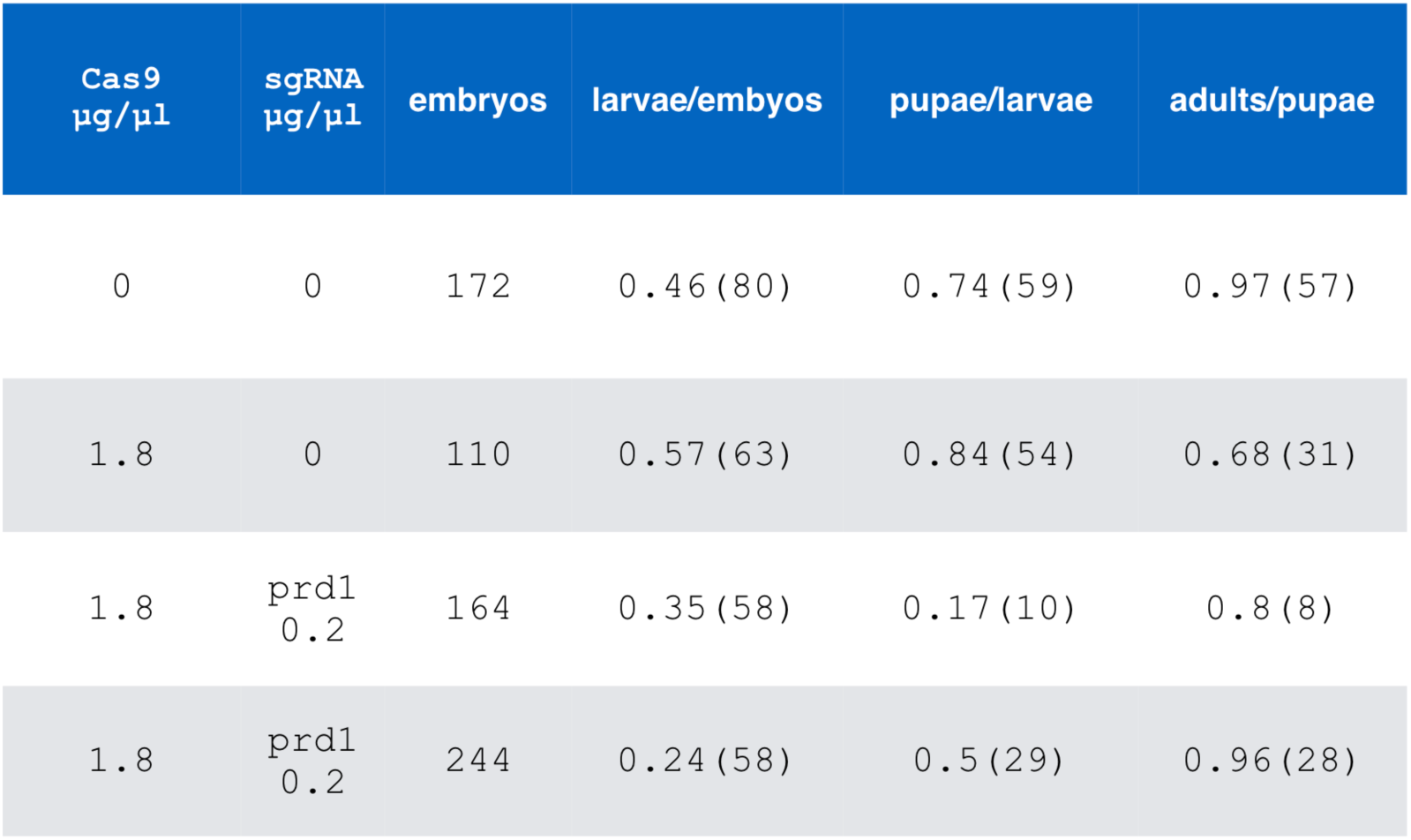
Survival rates after injecting of Ccprd-g1 and Ccprd-g2 RNPs into syncytial embryos. Numbers represent the percentage of survivors (actual number in parentheses) for a given specific developmental stage relative to number of individuals of the preceding stage.

| Cas9<br>µg/µl | sgRNA<br>µg/µl | embryos | larvae/embyos | pupae/larvae | adults/pupae |
| --- | --- | --- | --- | --- | --- |
| 0 | 0 | 172 | 0.46 (80) | 0.74 (59) | 0.97 (57) |
| 1.8 | 0 | 110 | 0.57 (63) | 0.84 (54) | 0.68 (31) |
| 1.8 | prd1<br>0.2 | 164 | 0.35 (58) | 0.17 (10) | 0.8 (8) |
| 1.8 | prd1<br>0.2 | 244 | 0.24 (58) | 0.5 (29) | 0.96 (28) |

